# Conformational space exploration of cryo-EM structures by variability refinement

**DOI:** 10.1101/2022.12.23.521827

**Authors:** Pavel V. Afonine, Alexia Gobet, Loïck Moissonnier, Billy K. Poon, Vincent Chaptal

## Abstract

Cryo-EM observation of biological samples enables visualization of sample heterogeneity, in the form of discrete states that are separatable, or continuous heterogeneity as a result of local protein motion before flash freezing. Variability analysis of this continuous heterogeneity describes the variance between a particle stack and a volume, and results in a map series describing the various steps undertaken by the sample in the particle stack. While this observation is absolutely stunning, it is very hard to pinpoint structural details to elements of the maps. In order to bridge the gap between observation and explanation, we designed a tool that refines an ensemble of structures into all the maps from variability analysis. Using this bundle of structures, it is easy to spot variable parts of the structure, as well as the parts that are not moving. Comparison with molecular dynamics simulations highlight the fact that the movements follow the same directions, albeit with different amplitudes. Ligand can also be investigated using this method. Variability refinement is available in the *Phenix* software suite, accessible under the program name *phenix*.*varref*.

## 1. Introduction

Proteins can be very mobile. They are subject to thermal entropy, and thus to deformations. Most of them undergo structural changes in line with their activity to fulfill their function. These movements range from local displacements to large domains movements.

Up until recently, these protein motions were mostly out of reach of high-resolution structures. X-ray crystallography was the main contributor of structures, and is by definition trying to diminish movements within the crystal to reach high resolution. Nevertheless, it was still possible to observe parts of the structure where the maps were poorly defined, implying local motion. The computation of Translation Libration Screw (TLS) parameters [1, 2] modeled large protein movements within the crystal which can be visualized as anisotropic displacement parameters. All in all, hints about protein motions could be deduced or calculated, but it was mostly linked to an absence of signal and always in the context of the crystal. Only solution NMR was able to capture protein dynamics, resulting in an ensemble of structure best fitting the distance restraints[3]. Unfortunately, this method is limited by the protein size, and solid-state NMR is still under development[4, 5] and cannot reach the large objects studied today.

Electron microscopy in cryogenic conditions (cryo-EM) made a real breakthrough in this regard by being able to visualize the sample under turnover conditions, and at high resolution[6, 7]. Cryo-EM is thus able to separate multiple conformations in the same sample. One potential application is in the elucidation of the structural changes during a catalytic cycle which will shed light on complex mechanisms. In addition to these large conformational changes, it is also possible to further dissect local proteins motions[8-12]. This is done by analyzing the variance of particle stacks to a volume along principal components, and outputs a series of volumes depicting protein motions.

These readily available local deformations are a real breakthrough in our understanding of protein movement dynamics. Many proteins use an energy source to undergo structural conformation, and defining energetic barriers is crucial to the fine understanding of their molecular mechanisms. The comparison of the maps constituting the movement is already a great achievement, it is undoubtably mesmerizing to observe the movements that were only conceptualized before. However, making sense of the motions requires detailed observation by a trained user, and it is hard to relate the motion to given parts of the protein.

In order to bridge the gap between observation and explanation, we designed a tool to create an ensemble of models fitting each map derived from variability analysis. The result is an ensemble of PDB models that can be used to describe protein movements, highlight transiently disordered parts of the structure, local rearrangements, helical twists and more. This ensemble of structures is complementary to high resolution structures, describes local motion and can serve as a base to more detailed dynamic studies. This tool has been made available in the Phenix software package[13] under the name *phenix*.*varref* and referred to as “*variability refinement*”.

## 2. Results

### 2.1. Generation of an ensemble of structures from a series of maps

Sample heterogeneity is observed in cryo-EM by fast freezing a sample under certain conditions, allowing local vibrations or larger structural changes to be preserved (Figure 1A). To reach high resolution, *in-silico* purification of a set of particles is required, yielding a high-resolution map (Figure 1B). This set of particles used to create the map still contains a certain level of heterogeneity (Figure 1C) that variability analysis can extract and output as a map series (Figure 1D), where each map differs a bit from each other relating a movement along a principal component axis[8].

**Figure 1:**
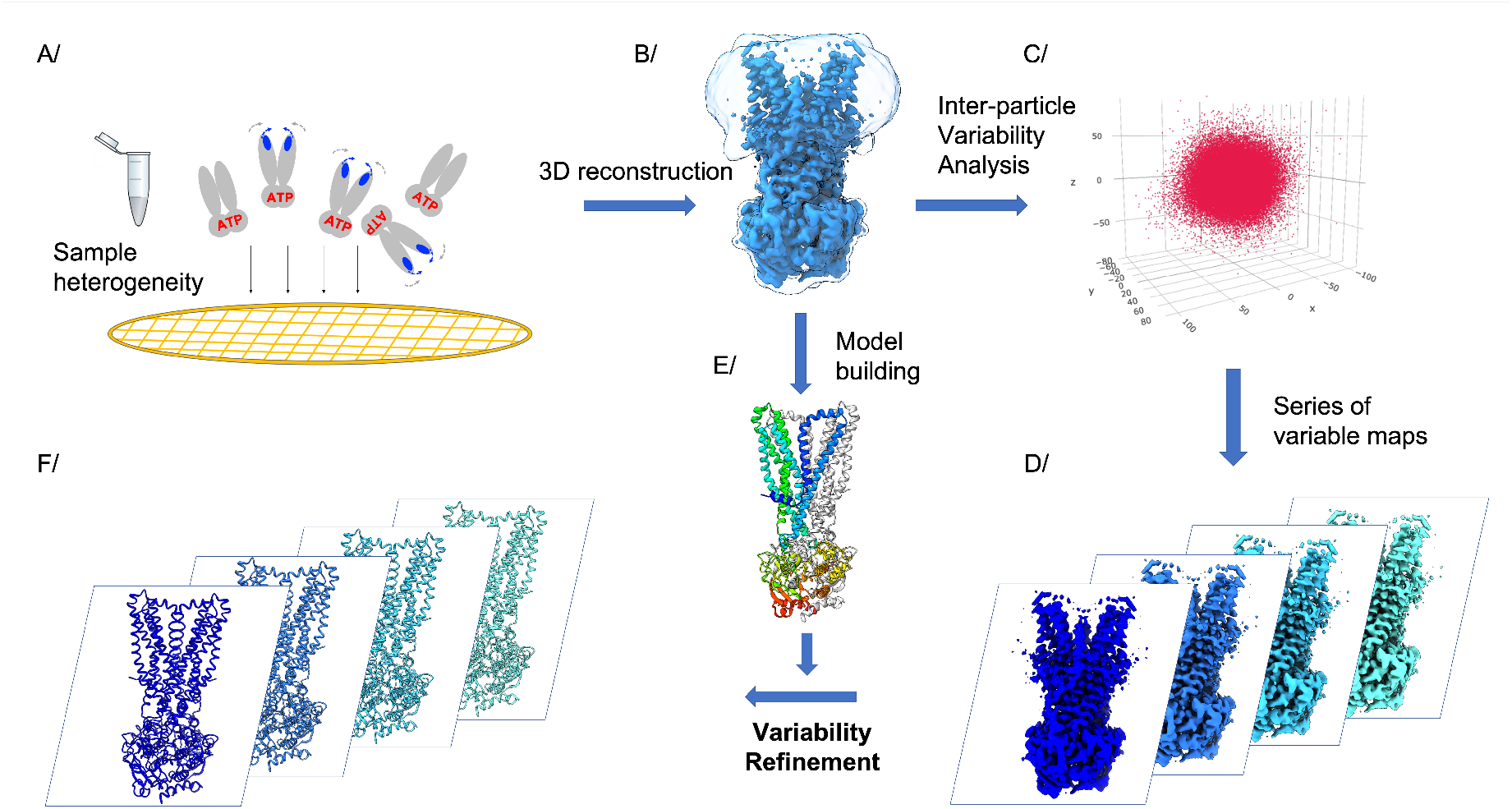
Schematic description of variability refinement. A/ sample heterogeneity is observed in cryo-EM by applying a sample on the grid and freezing its multiple states in liquid ethane. B/ 3D reconstruction from a set of particles yields an electrostatic potential map. C/ The 3D reconstruction is originating from a set of particles that differ more or less to the consensus. Variability can be observed using several software, here it is represented using 3DVA in CryoSPARC[17], where particles are separated by principal components axis. D/ from the variability analysis, a series of map is output. E/ a consensus model is built from the consensus map in B/. From the consensus model and the map series derived from 3DVA, variability refinement builds a model in each map to grasp the local movements.

In order to best describe these motions on the atomic level, a tool was created to create an ensemble of models, each model fitting its own map, given a high-resolution model. The procedure begins with pair-wise comparison of maps using correlation coefficient CC_box_[14] to sort and order maps by similarity (e.g., map i and map i+1 are more similar than map i and map i+2, and so on; map 1 and map M are most dissimilar; i=1,…M, M is the number of maps). Then the best matching map to the given starting atomic model is identified using map-model correlation coefficient CC_mask_[14]. This map is used to perform the first series of refinements. Each series of refinements against given map consists of N (N ∼10-100) refinements with identical settings but slightly perturbed input model. Performing the series of refinements rather than a single refinement is rationalized and illustrated in §3.2.4 in [15]. In a nutshell, this is done because refinement is a local optimization problem that has rather small convergence radius and is prone to be trapped in a nearest local minimum of the goal function being optimized. Starting with an ensemble of perturbed models allows for better sampling of the conformational space as well as for estimating an uncertainty in atomic positions[16]. The series of refinements against each map yields an ensemble of N refined models as well as the overall best model selected based in the CC_mask_. Next, the series N refinements proceeds with the following closest map and N refined models from the previous refinement as the starting point. In the end, the procedure generates NxM refined models and M overall best refined models.

### 2.2. Application of variability refinement to flexibility of the ABC transporter BmrA in the outward-facing conformation

ABC transporters are integral membrane proteins that harness the energy of ATP binding and hydrolysis to translocate substrates across biological membranes[18, 19]. BmrA is a multi-drug transporter from *B. subtilis*, that protects this microorganism in the competition for the biotope against *Streptomyces tendae*, which secretes cervimycin C as a biological weapon[20-22]. BmrA expels drugs out of the cell, thereby decreasing their concentration below cytotoxic levels, and thus protects the organism against xenobiotics. More generally, this mode of protection is found in every kingdom of life, and is a major mechanism leading to resistance to cancer treatments[23, 24], anti-fungal treatments[25-27] as well as antibiotic treatments[28, 29]. It is also found in all the physiological barriers in humans[30-32], having a key role in drug bio-availabilities throughout the organism.

The recent structure of BmrA in the outward-facing conformation, with and without substrate, in a pre-release state brought new insights to how the protein functions and to its energy landscape[33]. For instance, it was shown that no energy from ATP hydrolysis was needed to change the conformation from the inward-facing state to the outward-facing one, and that following substrate-release the transporter closes immediately to an occluded state. Plasticity and hydrophobic collapse were identified as the main drivers to a newly postulated swing mechanism of transport, allowing BmrA to handle a large variety of structurally unrelated drugs, a hallmark of multidrug transporter.

This outward-facing conformation is thought to be stable. Nevertheless, BmrA still undergoes local deformations that can be captured in the cryo-EM particle stack. 3D variability analysis of the high-resolution reconstruction shows movement of BmrA, reminiscent of how the transporter fluctuates around a central position (Figure 2-AB). Ensemble refinement in all the maps provided by 3DVA notably highlights the flexibility of the loop between TM1-2 (Figure 2C), which has been identified to be key in the protein’s adaptation to drugs[33]. The flexibility of this loop is detectable in the consensus map by judging the low contour of the density. This is due to the protein’s flexibility in this region which makes the loop hard to model. Ensemble refinement captures the flexibility, highlighted by variability analysis, of this zone.

**Figure 2:**
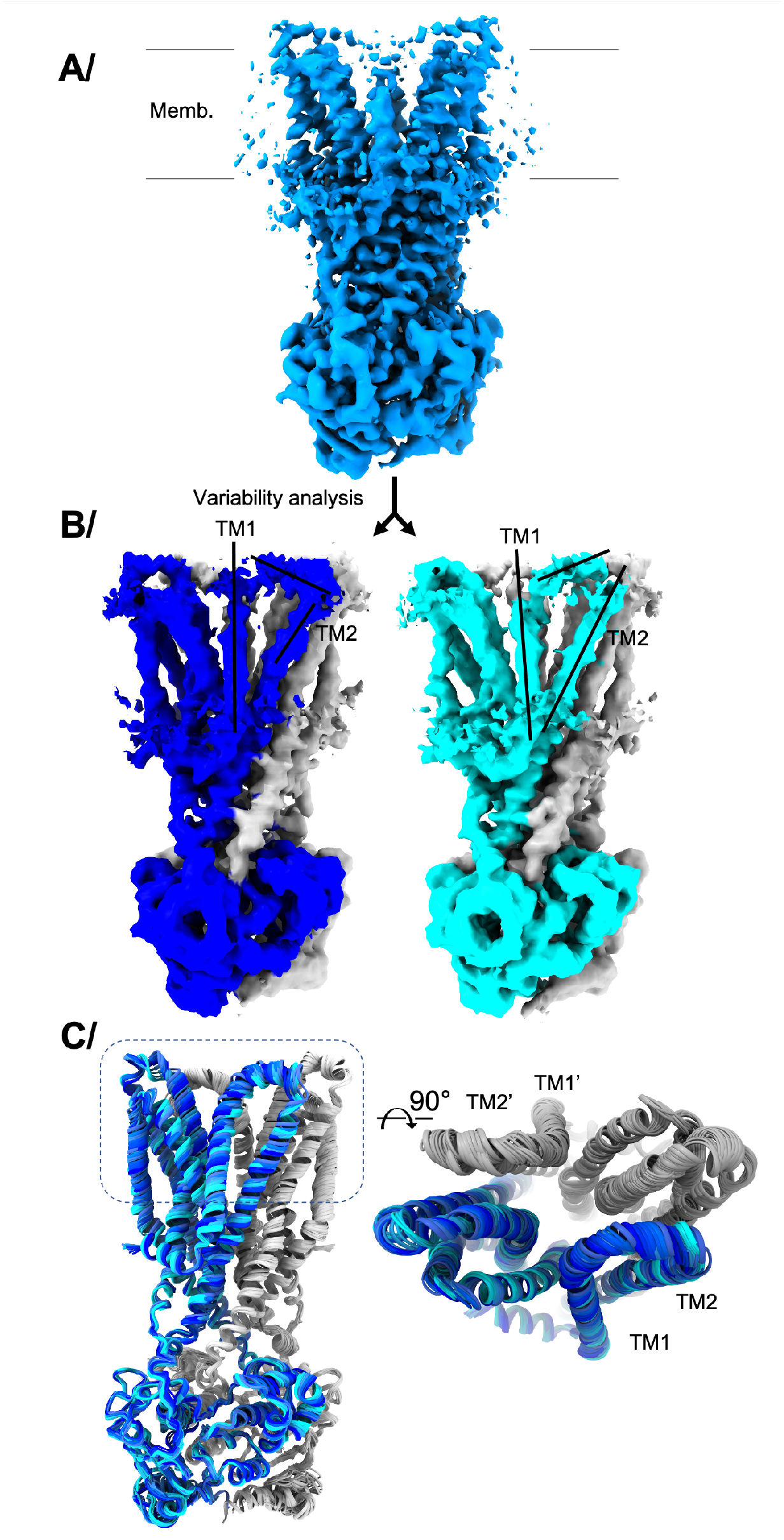
3D variability analysis of the ABC transporter BmrA in the outward-facing conformation. **A/** Consensus reconstruction shown perpendicular to the membrane plane. **B/** Variability analysis with the two outmost maps displayed. One monomer shown in blue and the other in silver. Movements range from dark blue to cyan. Different positions of TM1, TM2 and the loop between the two are displayed with black bars. **C/** Variability refinement of BmrA in all the maps originating from variability analysis. The protein is shown in cartoon, with one monomer in silver and the other in blue, with a color gradient ranging from dark blue to cyan according to B/. 20 models are displayed to visualize all the positions sampled by the protein. The overall rmsd is 0.6 Å / 1065 residues (dimer), with local displacements in TM1-2 and other flexible regions of 2 Å per residue.

### 2.3. Comparison of movements between variability refinement and molecular dynamics simulations

All-atom molecular dynamics (MD) simulation of BmrA in a phospholipid bilayer has been performed in the same conditions as the structure[33], thus allowing the comparison of the observed movements (Figure 3). The first observation is that the magnitude of the movements is a bit larger in the MD simulation compared to the variability contained within the particle stack used to recreate the volume at high resolution, which is in accordance with the expectation that restrained coordinates are required to reach high resolution (cryo-EM map) compared to giving freedom to the system to explore conformational space (MD simulation). Upon closer look, it becomes clear that the movements revealed by variability refinement correspond to the same ones observed by MD simulation. The larger movements of the TM1-2 loop, as well as the TM3-4 and TM5-6 linkers, are clearly distinguishable in the bundle of structures and correspond to the most variable regions of the transporter in this conformation.

**Figure 3:**
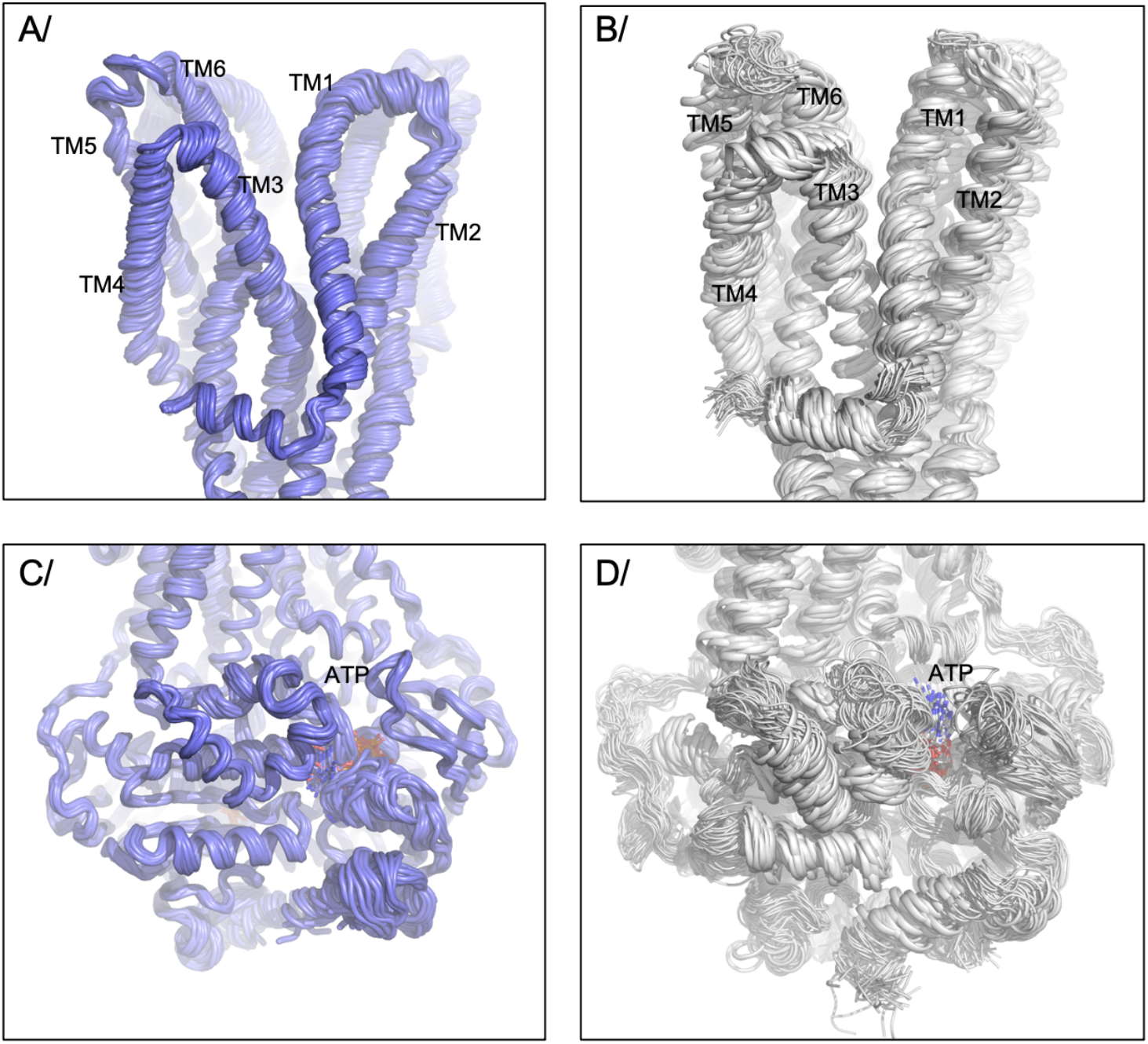
Comparison of variable parts of the structure compared to molecular dynamics simulation. Panel A/ and C/ show the movement derived from variability refinement, while B and D show movements undergone during molecular dynamics simulation. The movement are separated for trans membrane domain on top with trans-membrane helices labeled, and the nucleotide binding domain at the bottom with ATP labeled. Protein shown in cartoon and ATP in sticks colored by atom type. In the MD simulation, the rmsd between models varies between 1.6 and 2.7 Å for 590 residues (monomer).

In the nucleotide-binding domain (NBD), the vibration observed in MD simulations is also transcribed in the bundle of structures created by variability refinement, even though it is clear that this region is moving less than the trans-membrane region. Ligand movement is also seen, and detailed below.

Overall, the structure bundle is revealing the regions of the protein undergoing structural rearrangements. Even though the movement amplitude is a bit less in this particle stack compared to the MD simulation, it is striking to notice that the movements described for the protein are in the same direction for each secondary structure element. The structure bundle created by variability refinement grants this visualization, as a first approximation to larger protein movements.

### 2.4. Movements of ligands

Ligands are also handled by variability refinement. Note that an important strategy for variability analysis is the calculation at lower resolution than the one used to obtain the highest resolution map since protein deformations are more likely to be seen at lower resolution. Thus, ligands density can become less clear or become lost, so caution is advertised to interpret ligand movements using this method. If needed, restraints can be provided to link the ligand to the protein so it can ride along with the deformations. This can be the case for ions for instance, while larger ligands will be less subjected to density loss for the entire length of the ligand. In the structure of BmrA, ATP has a well-defined density in the consensus map, and while density is still visible in the 3DVA analysis, its quality is affected by the lower resolution of the calculation. Thus, it is important to add distance restraints otherwise the harsh procedure to escape local energy minimum will trigger the protein loops to move into ligand density, leading to over-refined structures, and doubtful results. Comparison of ATP in the MD simulation with the structure bundle coming from variability refinement (Figure 4A) gives hints on the movements undergone by the ligand within the structure. Figure 4A highlights that ATP can undergo significant movements in the outward-facing conformation, in the two types of studies. Molecular details shown in Figure 4B-D follow the movements observed for the whole transporter (Figure 3), where the amplitude of movements is larger in the MD simulation. Overall, this allows to visualize how the ligand move in concert with the protein, and follows protein movements. Variability analysis and refinement thus helps to understand local movements within the structure.

**Figure 4:**
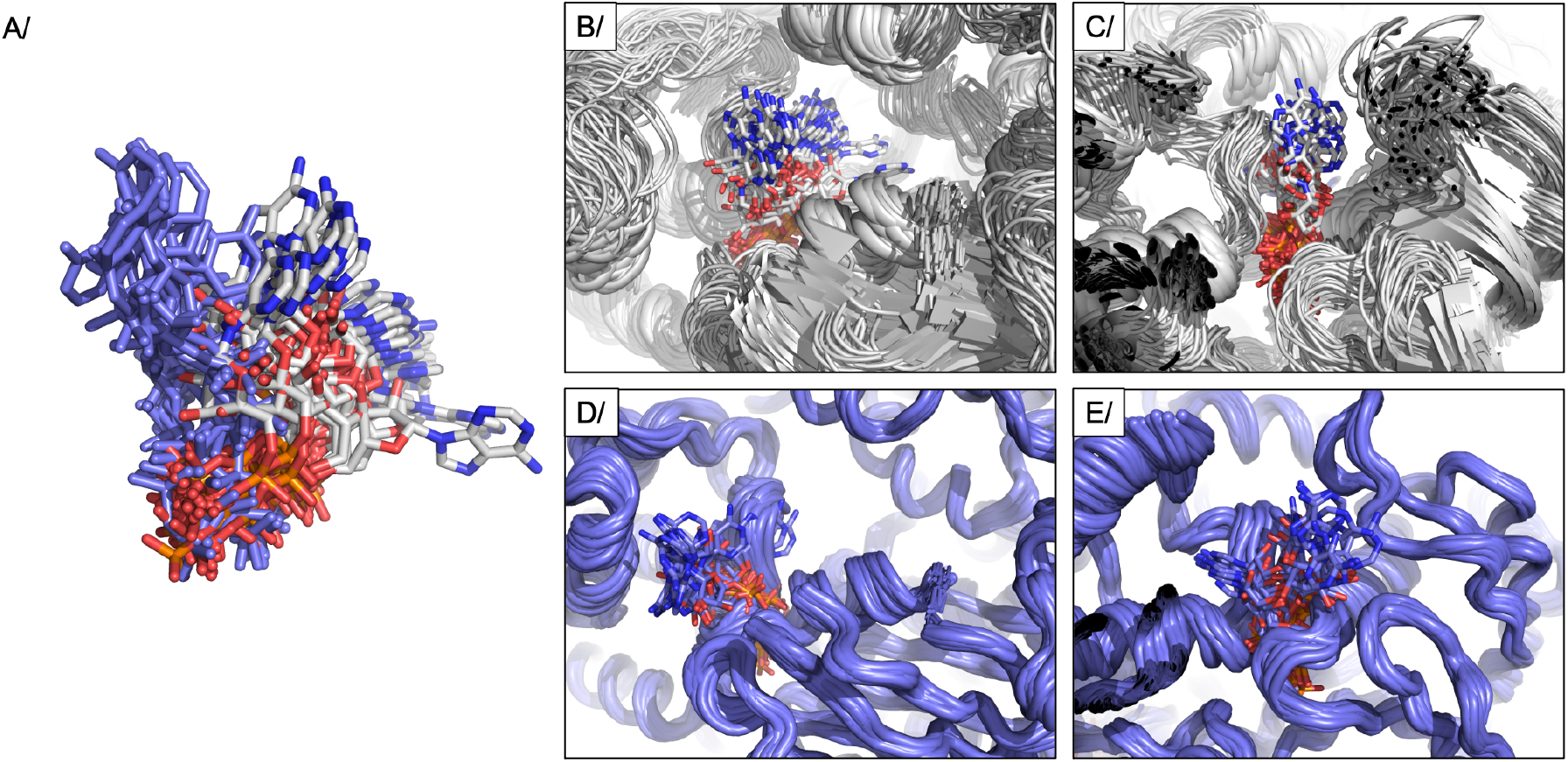
Comparison of ATP movement using variability refinement and molecular dynamics simulations, in the same conditions. A/ ATP overlay between molecular dynamics simulations (colored by atom type) and variability refinement (Blue). B/ and C/ show 2 views of the ATP-binding site from molecular dynamics simulations. C/ and D/ show the same views for the ATP binding-site from variability refinement.

### 2.5. Application of phenix.varref to large complexes

Variability refinement was tested on a large super-complex to assess the robustness of the software. CryoEM maps derived from variability analysis of EMPIAR-10180 entry were graciously shared by the CryoSPARC team for this analysis. This structure of the pre-catalytic yeast spliceosome[34] contains 39 protein chains and 5 nucleic acid chains, for a total of 2.5 MDa. Variability refinement successfully modeled each chain in the map, faithfully following the movement observed in cryoEM maps (Figure 5AB). Each chain undergoes different movement according to the deformation observed in the cryoEM structure, allowing for investigators to observe protein movements within the structure. In addition to being a large complex, this structure is also of modest resolution (7 Å overall), making that variability analysis was calculated at 8 Å resolution. There is thus a lack of details to guide the exact reconstruction of the super-complex, and this is especially notable in some regions of the complex like exemplified on the top-right part of the structure (Figure 5A), for which electron maps show the least quality, synonym of maximum displacement. The placement of a model in this context is a major challenge, and care is advertised in the interpretation of movements in this region. Undeniably, a higher resolution version of this structure will render reconstructions more accurate, and a better description of movements going on in this region of the complex.

**Figure 5:**
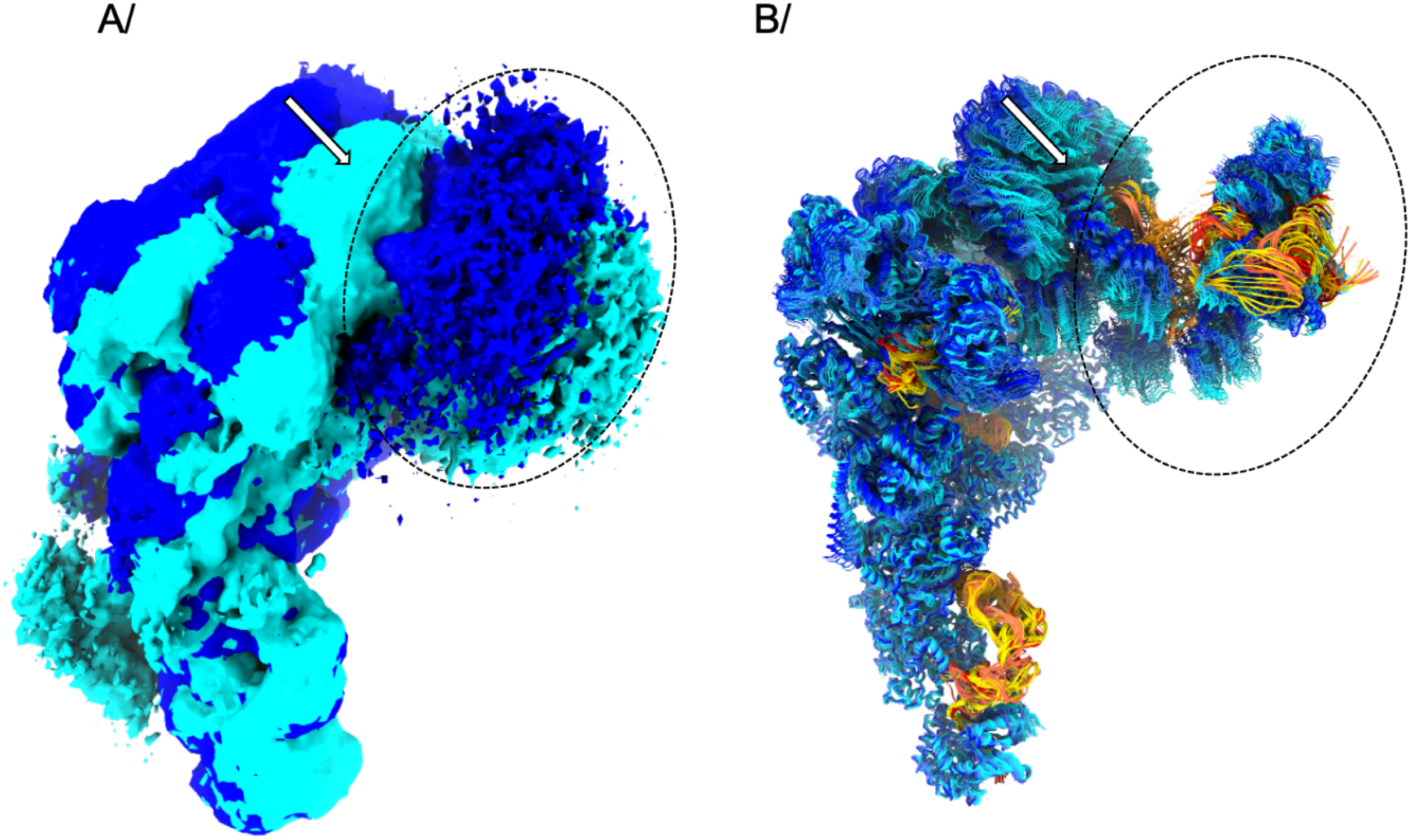
Variability refinement of the spliceosome (EMPIAR-10180). A/ Variability analysis of EMPIAR-10180 calculated by CryoSPARC, with the two outmost maps displayed; movements range from dark blue to cyan. B/ Variability refinement of the spliceosome, with protein chains colored from dark blue to cyan (following the maps coloring), and DNA chains from red to yellow. The white arrow exemplifies a movement as seen in the cryoEM maps (A/), and its calculation using variability refinement (B/). The dashed circle points to an area where the signal is very low and thus the model generated has to be interpreted with caution.

## 3. Discussion/ Conclusion

An interesting feature of cryo-EM is the ability to observe a sample in multiple states. These states can represent the variability inherent in the sample and various methods exist to analyze the variability between a volume and a set of particles. The result is a map series that recreates a movement. While this movement is a wonderful way to visualize how a protein deforms around a given conformation, it is hard to dissect the movement and pinpoint local deformations on a map. We created the tool *variability refinement* to refine an ensemble of structures for each map originating from variability analysis, thus creating a bundle of structures that describes the movement. It is then easy to spot the variable parts of the protein, and to define the amino-acids involved in the movement and those that are not. Investigators can have access to key deformation hot spots, and measure and quantify the movements using standard PDB tools that are widely available.

It would be interesting to add time to the structural changes observed. Admittedly, the movement output by variability analysis is a principal component analysis of particle variability, and the linearity of the movement is originating from computation[8]. Nevertheless, the ensemble of maps, and now of structures, depicts protein variability in this experimental condition, and in this regard, the movie of the protein moving along the principal component represents a plausible event. By combining movements along several principal components, a comprehensive view of the movement can be grasped. Therefore, considering the set of maps as a trajectory is a believable proxy. Comparison with MD simulations reveals that the direction of the movements observed for the ABC transporter BmrA in the outward-facing conformation are the same, and that the flexible regions correspond between MD simulation and variability analysis. The movement amplitude is however smaller in this variability analysis, revealing that the time that the protein would use to undergo these conformational changes is smaller than the MD simulation time, of 500ns in this case. Nevertheless, it is interesting to note that the movements described by both methods follow the same directions, giving hints to the deformable parts within the structure.

The variability analysis of BmrA in this outward-facing conformation is in the small range, corresponding to local displacements. BmrA also undergoes significantly larger movements in the inward-facing conformation, currently being investigated and tested for variability refinement but not shown here as the analysis is still preliminary. We also showed that variability refinement can be used for even larger systems like the spliceosome[34] that undergo much larger structural changes with movements of large domains, and it will be interesting to test it for other similar systems [35, 36]. The *variability refinement* tool is publicly available in Phenix starting with the version dev-4799 for investigators to use.

Now that Artificial Intelligence (AI) has undergone a major leap in deciphering the code to predict protein structures from sequence[37], the next step is to know how these proteins move. Using variability refinement, the community has now a method to create a new type of learning set for AI to predict protein movements, and steps will be undertaken to make this data publicly available. Along the same lines, the very recent release of another type of movement descriptor by 3DFLEX [38] allows for a whole visualization of the movement, not limited to linear interpretation, and reflecting conformational flexibility. Steps are being undertaken to adapt variability refinement to this new calculation of protein flexibility.

## 4. Material and methods

### 4.1. Generation of the map ensemble for the ABC transporter BmrA

3D variability analysis for BmrA is performed in CryoSPARC v3.3. 3D variability analysis was carried on 327,764 particles used for the consensus high-resolution reconstruction [33] (PDB: 7OW8. EMDB: 13095). Variability was computed in simple mode, at 6 Å resolution and C1 symmetry.

### 4.2. Variability refinement of BmrA

Maps originating from 3DVA were input to *phenix.varref* along with the refined structure and a resolution of 6 Å, on a 72-core Linux calculation server (Intel® Xeon® 6240 at 2.60 GHz). Using 50 cores, the job ran in about 6 hours. Some distance restraints were used between ATP atoms and some residues in the neighboring loops.

### 4.3. Variability refinement of the spliceosome

Maps originating from 3DVA were generously shared by the cryoSPARC team, performed on EMPIAR entry 10180. Only the component 0 was used to assess the feasibility of using *phenix.varref* on such a large super-complex. 3DVA was calculated by the cryoSPARC team using the particles and mask from the EMPIAR entry, using a filter resolution of 8 Å. Finally, maps were displayed with a filter resolution of 6 Å. Using 50 cores (on an Intel® Xeon® 6240 at 2.60 GHz), the job ran in about 6 days.

### 4.4. Usage of phenix.varref

Variability refinement in phenix can be launched using the command-line: phenix.varref name.pdb frame_*.mrc resolution=4 nproc=50 models_per_map=50 where name.pdb is the refined model, frame_*.mrc points to the maps from variability refinement, the resolution needs to be specified, nproc is the number of processors, and models_per_map indicate the initial number of models created, 50 by default.

Figures were created with ChimeraX[39] and Pymol[40].

## Author contributions

VC initiated the study. PVA and BKP wrote the software. AG, LM and VC tested the program on their data. All authors wrote the manuscript.

## Acknowledgments

This work was supported by the CNRS, and the French National Research Agency grant number ANR-19-CE11-0023-01 to VC and AG. VC wishes to thank Juliette Martin for sharing the molecular dynamics data, and Xavier Robert for his help running the crystallography calculation cluster. PVA and BKP thanks the NIH (grants R01GM071939, P01GM063210 and R24GM141254) and the PHENIX Industrial Consortium for support of the PHENIX project, as well as the support by the US Department of Energy under Contract No. DE-AC02-05CH11231. The authors thank the cryoSPARC team for sharing the spliceosome 3DVA maps.

